# Modulation of the spontaneous hemodynamic response function across levels of consciousness

**DOI:** 10.1101/401547

**Authors:** Guo-Rong Wu, Carol Di Perri, Vanessa Charland-Verville, Charlotte Martial, Steven Laureys, Daniele Marinazzo

**Affiliations:** Department of Data Analysis, University of Ghent, B9000 Ghent, Belgium; Key Laboratory of Cognition and Personality, Faculty of Psychology, Southwest University, Chongqing 400715, China; Coma Science Group, GIGA Research Center, University of Liège, B4000 Liège, Belgium; Center for Clinical Brain Sciences, Centre for Dementia Prevention, UK Dementia Research Institute, University of Edinburgh, EH16 4SB Edinburgh, United Kingdom

## Abstract

Functional imaging research has already contributed with several results to the study of neural correlates of consciousness. Apart from task-related activation derived in fMRI, PET based glucose metabolism rate or cerebral blood flow account for a considerable proportion of the study of brain activity under different levels of consciousness. Resting state functional connectivity MRI is playing a crucial role to explore the consciousness related functional integration, successfully complementing PET, another widely used neuroimaging technique. Here, spontaneous hemodynamic response is introduced to characterize resting state brain activity giving information on the local metabolism (neurovascular coupling), and useful to improve the time-resolved activity and connectivity measures based on BOLD fMRI. This voxel-wise measure is then used to investigate the loss of consciousness under Propofol anesthesia and unresponsive wakefulness syndrome. The dysfunction of hemodynamic response in precuneus and posterior cingulate is found to be a common principle underlying loss of consciousness in both conditions. The thalamus appears to be less obviously modulated by Propofol, compared with frontoparietal regions. However, a significant increase in spontaneous thalamic hemodynamic response was found in patients in unresponsive wakefulness syndrome compared with healthy control. Our results ultimately show that anesthesia- or pathology-induced neurovascular coupling could be tracked by modulated spontaneous hemodynamic response derived from resting state fMRI.

## Introduction

In order to provide effective assistance to diagnosis, prognosis, and assess potential treatments, it is crucial to understand the physiological basis of consciousness in pathological or pharmacological coma. Advanced neuroimaging techniques have significantly expanded our knowledge of neural correlates of consciousness level in human brain. For instance, FDG-PET imaging and fMRI have shown that altered metabolism and connectivity in thalamus, frontoparietal and default mode networks (Laureys, 2005; Laureys et al., 2000a; Laureys et al., 2000b; Laureys et al., 2002; Laureys et al., 2004) are found in patients with disorders of consciousness; although electrophysiological techniques have discovered some covert rhythm signs of consciousness (Gugino et al., 2001; Vijayan et al., 2013). Finding high test-retest reliable evidences within or across multimodal imaging are a necessary step to obtain clinically applicable neural markers of consciousness. This kind of work has gained extensive attention, and not confined to consciousness, especially in fMRI studies (Plichta et al., 2012). A clinical validation study suggested that PET imaging could have a higher diagnostic precision than event-related fMRI in disorders of consciousness (Slender et al., 2014). This call for an improved mining of the dynamic information by means of advanced fMRI models, considering the superior spatiotemporal resolution of fMRI.

Consciousness has two major components: awareness of environment and of self (i.e. the content of consciousness), and wakefulness (i.e. the level of consciousness) (Laureys, 2005). The vegetative state/unresponsive wakefulness syndrome (UWS) is the most dramatic model of dissociation between wakefulness and awareness. Indeed, patients in unresponsive wakefulness syndrome are awake but seemingly not aware. In the last decade neuroimaging studies on these patients have extensively investigated the neural correlates of awareness, mainly exploring the correlation between awareness and global brain function (Laureys et al., 1999b), regional brain function, brain activation induced by passive external stimulation (Laureys and Schiff, 2012) or mental imagery task (Owen et al., 2005), and changes in resting state connectivity (Laureys et al., 2000b; Laureys et al., 1999a; Laureys et al., 1999b). Similar approaches have also been applied to identify the neural correlates: of different states of consciousness as induced by anesthetic drugs manipulation (Boveroux et al., 2010; DiFrancesco et al., 2013; Hudetz, 2012). Compared to global brain metabolism, regional metabolic dysfunction is a more reliable maker of individual conscious-unconscious state, transition (Laureys, 2005). Studies of passive stimuli have shown promising results, especially for minimally conscious state patients, but the neuronal responses cannot be always stable (Laureys and Schiff, 2012); as a consequence inference of cognitive function from these cerebral activations is controversial (Menon et al., 1999; Schiff and Plum, 1999). Resting state functional connectivity (FC) plays a critical role in the quantification of brain function integration sustaining the state of consciousness (Boveroux et al., 2010; Laureys et al., 2000b; Laureys et al., 1999a; Laureys et al., 1999b). Although it well confirmed previous FDG-PET findings (Laureys and Schiff, 2012), as now this approach is susceptible to some limitations, such as the neurovascular connection anatomy (Tak et al., 2014) and motion artifacts (Power et al., 2012), which may contribute to high proportion of the resting state FC signal commonly detected by BOLD fMRI.

BOLD-fMRI hemodynamic response describes the vascular oxygenation changes following a neuronal impulse response and is in this sense a more direct measure of neural activity than PET. Here we propose another possible marker of cerebral metabolic patterns of altered levels of consciousness: we investigate the shape pattern of the hemodynamic response using resting state fMRI data in UWS patients and healthy subjects under Propofol sedation. The hemodynamic response correlates of consciousness will be explored and compared between UWS and anesthesia. Apart from being a local biomarker, the hemodynamic response affects activation and connectivity (Rangaprakash et al., 2018). A subject- and voxel-vise estimation of its shape would allow a better estimate of these latter measures. Previous PET (Boly et al., 2011; Laureys, 2005; Laureys et al., 2004) and fMRI studies (Guldenmund et al., 2013) reported that thalamus, fronto-parietal cortical areas, salience network and default mode network (DMN) exhibit state-dependent spontaneous hemodynamic response in the resting state. We will here aim to verify if these local modulations are replicated with the newly proposed measure.

## Materials and Methods

### Subjects

Two datasets, the Propofol and the UWS one were used in this study. Twenty-one healthy right-handed volunteers were included in the Propofol dataset. Twenty UWS patients and thirty-two healthy controls (HC) participated in the UWS study. Diagnosis of UWS was made after repeated behavioral assessments by experienced and trained neuropsychologists using the Coma Recovery Scale-Revised (CRS-R; (Giacino et al., 2004; Wannez et al., 2017)). Written informed consent to participate in the study was obtained from the healthy subjects and from the legal representatives of the patients. None of the healthy subjects had a history of head trauma or surgery, mental illness, drug addiction, asthma, motion sickness, or previous problems during anesthesia. The study was approved by the Ethics Committee of the Medical School of the University of Liège (University Hospital, Liège, Belgium). The relevant demographic and clinical information can be found in Tattle 1 and Table 2. For the UWS dataset, no statistically significant differences in age and sex were found between the control group and the patients’ one.

**Table 1:** Demographics test for all subjects, a: Chi-Square goodness of fit test; b: Pearson’s Chi-square test for independence.

|  | UWS (n=13) | Healthy Controls (n=25) | Propofol (n=21) |
| --- | --- | --- | --- |
| Age (y) | Mean=42.8 | Mean=42.8 | Mean=23.4 |
|  | SD=16.5 | SD=15.7 | SD=4.2 |
| | $t(36)=0.0011, p = 0.9991$ | | |
| Sex | M=7 | M=16 | M=5 |
|  | F=6 | F=9 | F=16 |
| | $\chi^2(1)= 0.08$ | $\chi^2(1)= 1.96$ | $\chi^2(1)= 5.76$ |
| | $p=0.78^a$ | $p=0.16^a$ | $p= 0.02$ |
| | $\chi^2(1)= 0.37, p=0.54^b$ | | |

**Table 2:** Demographics and accident etiology for IJWS patients

| Patient | Sex | Age (y) | Accident Etiology |
| --- | --- | --- | --- |
| sub01 | F | 44 | anoxia |
| sub02 | M | 15 | TBI |
| sub03 | M | 54 | post meningitis a<br>pneumococcus |
| sub04 | F | 38 | anoxia |
| sub05 | M | 33 | intracranial hemorrhage |
| sub06 | M | 28 | mixed TBI-anoxia |
| sub07 | M | 43 | anoxia |
| sub08 | F | 69 | anoxia |
| sub09 | F | 57 | TBI |
| sub10 | M | 73 | TBI |
| sub11 | F | 31 | TBI |
| sub12 | F | 36 | anoxia |
| sub13 | M | 36 | anoxia |

### Functional Data Acquisition

The Propofol dataset considered in current study has already been published in (Boveroux et al., 2010). Functional MRI acquisition consisted of resting-state fMRI volumes repeated in four clinical states only for 21 healthy volunteers: normal wakefulness (Wl), mild sedation (Si), deep sedation (S2), and recovery of consciousness (W2). The temporal order of mild- and deep-sedation conditions was randomized. The typical scan duration was half an hour in each condition. The number of scans per session was matched in each subject to obtain a similar number of scans in all four clinical states (mean±SD, 251 ± 77 scans/session).

There is only one session of resting state fMRI for each subject in the UWS dataset, each one contains 300 scans.

All functional images were acquired on a 3 Tesla Siemens Allegra scanner (Siemens AG, Munich, Germany);

Propofol dataset: Echo Planar Imaging sequence using 32 slices; repetition time (TR)=2460ms, echo time=40ms, field of view = 220mm, voxel size=3.45×3.45×3mm, and matrix size=64×64 ×32).

UWS dataset: Echo Planar Imaging sequence using 32 slices; repetition time (TR)=2000ms, echo time=30ms, field of view = 384mm, voxel size=3.44×3.44×3mm, and matrix size=64× 64×32).

### Data Preprocessing

All structural images in both datasets were manually reoriented to the anterior commissure and segmented into grey matter, white matter (WM), cerebrospinal fluid (CSF), skull, and soft tissue outside the brain, using the standard segmentation option in SPM 12. Then a study-based template was created, in order to minimize deformity due to atrophic brains.

Resting-state fMRI data preprocessing was subsequently carried out using both AFNI and SPM12 packages. First, the EPI volumes were corrected for the temporal difference in acquisition among different slices, and then the images were realigned to the first volume for head-motion correction. 8 UWS patients and 7 healthy control subjects were excluded from the. dataset because either translation or rotation exceeded ±1.5 mm or ±1.5°, or mean framewise displacement (FD) exceeded 0.5, resulting in 13 UWS patient and 25 healthy controls which were used in the analysis, and 22 sessions in Propofol group subjects were also excluded. The resulting volumes were then despiked using AFNI’s 3dDespike algorithm to mitigate the impact of outliers. The mean BOLD image across all realigned volumes was coregistered with the structural image, and the resulting warps applied to all the despiked BOLD volumes. Finally all the coregistered BOLD images were smoothed (8 mm full-width half-maximum) and spatially normalized into MNI space.

Several parameters were included in a linear regression to remove possible spurious variances from the data. These were i) six head motion parameters obtained in the realigning step, ii) non-neuronal sources of noise estimated using the anatomical component correction method (aCompCor, the representative signals of no intei’est from subject-specific, WM and CSF included the top five principal components from WM and the top five from CSF mask) (Behzadi et al., 2007). Then the residual time series were linearly detrended and temporally band-pass filtered (0.008–0.1 Hz).

### Spontaneous point process event and HRF retrieval

We employed a blind hemodynamic response function (HRF) retrieval technique specially developed for resting-state BOLD-fMRI signal (Wu et al., 2013), which considers the resting-state BOLD signal as driven by spontaneous point process events. A linear time-invariant model for the observed resting state BOLD response is assumed (Boynton et al., 1996; Dale and Buckner, 1997). We consider a common HRF is shared across the various spontaneous point process events at a given voxel, to ensure a more robust estimation. The BOLD-fMRI signal *y*(*t*) at a particular voxel is given by:

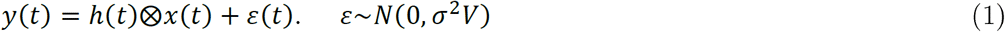

where *x*(*t*) is the sum of time-shifted delta functions, centered at the onset of each spontaneous point process event and *h(t)* is the (unknown) hemodynamic response to these events, *ε*(*t*) represents additive noise and ⊗ denotes convolution. The noise errors are not independent in time due to aliased biorhythms and unmodellecl neural activity, and are accounted for using an AR(p) model during the parameter estimation (we set p=2 in current study). In practice, *y*(*t*) is not sampled continuously in time, but rather at discrete intervals, i.e. TR. Consequently, the convolution was performed in TR temporal resolution in a previous study (Wu et al., 2013). Given that spontaneous point process event onsets do not need to be synchronized with scans, here we perform the convolution at a higher temporal resolution with N time points per scan then down-sampled to TR temporal resolution at Equation (2). Although no explicit external inputs exist in resting-state fMRI acquisitions, we still could retrieve the timing of these spontaneous events by blind deconvolution technique (Wu et al., 2013). The peak of BOLD response lags behind the peak of neural activation is presumed to 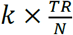 seconds (where 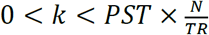, peristimulus time, PST). The timing set *S* of these resting-state BOLD transients is defined as the time points exceeding a given threshold around a local peak, can be detected by the following way: *S*{*i*} *= t*_*i*_, *y(t*_*i*_) ≥ *m &.y(t*_*i*_*) ≥ y*(*t*_*i*_ ± τ), where we set τ = 1,2 and *m = μ + σ* (i.e. SD) in current study. While the exact time lags can be obtained by minimizing the mean squared error of equation (1), i.e. the optimization problem:

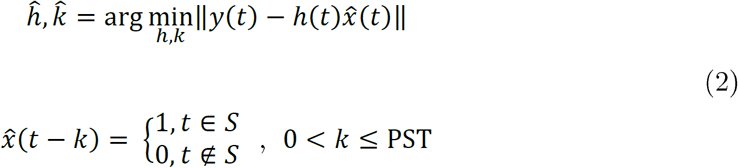

In order to escape motion artifacts induced pseudo point process events, a temporalmask with FD<0.3 was added to exclude these bad pseudo-event onsets from timing set *S* by means of data scrubbing (Power et al., 2012).

The method described above makes no assumptions about the exact shape or functional form of the hemodynamic responses. The application of prior knowledge about possible hemodynamic response shapes could reduce the bias in the linear estimation framework especially for the low signal noise ratio dataset, and sharply reduce the computational cost. Here we assume that the hemodynamic response for all resting state spontaneous point process events and at all locations in the brain are fully contained in a *d*-dimensional linear sub-space *H* of ℜ^d^, then, any hemodynamic response *h* can be represented uniquely as the linear combination of the corresponding basis vectors. The canonical HRF in SPM with its delay and dispersion derivatives are employed as the basis functions in current study (Friston et al., 1998) (we denote it as *canon2dd* model).

To characterize the shape of hemodynamic response, three parameters of the HRF, namely response height, time to peak, Full Width at Half Maximum (FWHM), were estimated, which could be interpretable in terms of potential measures for response magnitude, latency and duration of neuronal activity (Lindquist and Wager, 2007). In Figure 1 some results are summarized, evidencing the BOLD point processes with the pseudo-events generating them.

**Figure 1.**
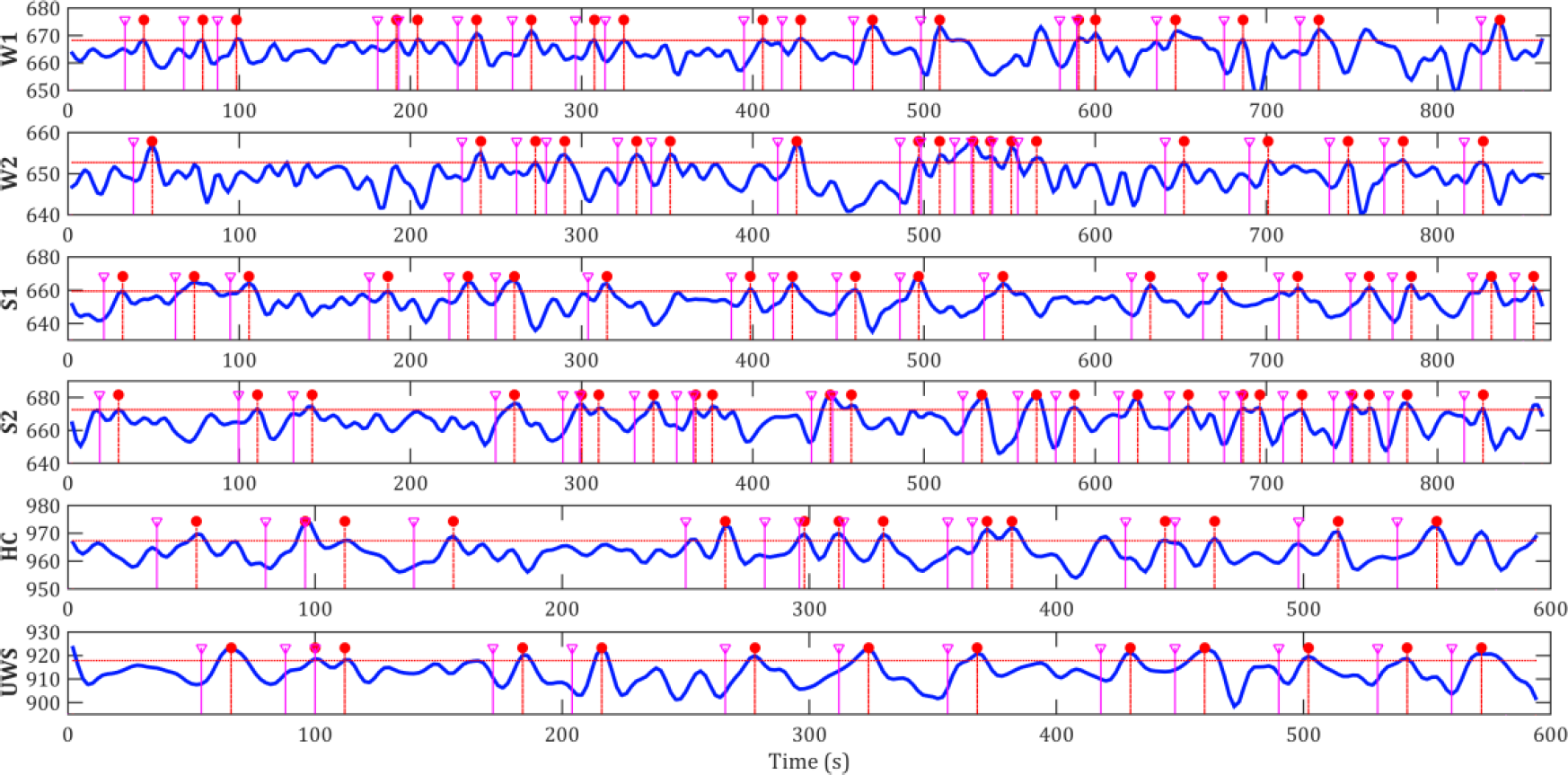
Retrieval of point processes and pseudo-event times: the red horizontal dashed line are the threshold (mean+SD), the red vertical dashed lines represent the point process, the purple lines indicate the pseudo neural events onset times (lagging behind the BOLD point process). Each signal shown here comes from one voxel in the precuneus, the first four rows represent the same voxel for the same subject. Due to motion some above-threshold time points have been “scrubbed”.

The toolbox to perform these analyses (in Matlab, Python, Docker Container, and BIDS app) is freely available on the NITRC page https://www.nitrc.org/proiects/rshrf.

### Statistical Analysis

We retrieved resting state HRF for all the voxels contained within the cerebrum using the AAL template (Tzourio-Mazoyer et al., 2002). HRF parameters for each subject were entered into a random-effects analysis (one-way AN OVA) within subjects, with three covariates (age, sex and mean FD) to identify regions which showed significant activity differences among four clinical states, a linear t contrast was computed, searching for a linear relationship between HRF and the level of consciousness across the four conditions (contrast (Wl. W2, SI, S2): [1.5 0.5 −1.5 −0.5]) (Boveroux et al., 2010). A pairwise t-test between W1 and S2 (the two extreme states) was further performed to compare with the results from the UWS dataset.

A two-sample t-test with three covariates (age, sex and mean FD (Power et al., 2012)) was implemented to map group difference of HRF parameters between control subjects and UWS patients.

The consistent group difference maps from the two different datasets were obtained using conjunctions (minimum statistic compared to the conjunction null) (Nichols et al., 2005).

Type I error due to multiple comparisons across voxels was controlled by false discovery rate (Chumbley et al., 2010). Statistical significance for group analysis was set at P_FDR_<0.05, derived from Gaussian random field theory.

## Results

### Spatial distributions of resting state HRF parameters

The temporal interval of spontaneous point process events is inhomogeneously distributed (Figure 2), a feature that enables a more robust GLM estimation for HRF retrieval. HRF parameters of each voxel are estimated and mapped on the brain (Figure 3). The median maps of each HRF parameters exhibit spatially heterogeneity across different level of consciousness and HRF models (Figure 3). Similardistributions are present in Wl, W2, and control group, higher response height and FWHM appear in frontal lobe and precuneus.

**Figure 2.**
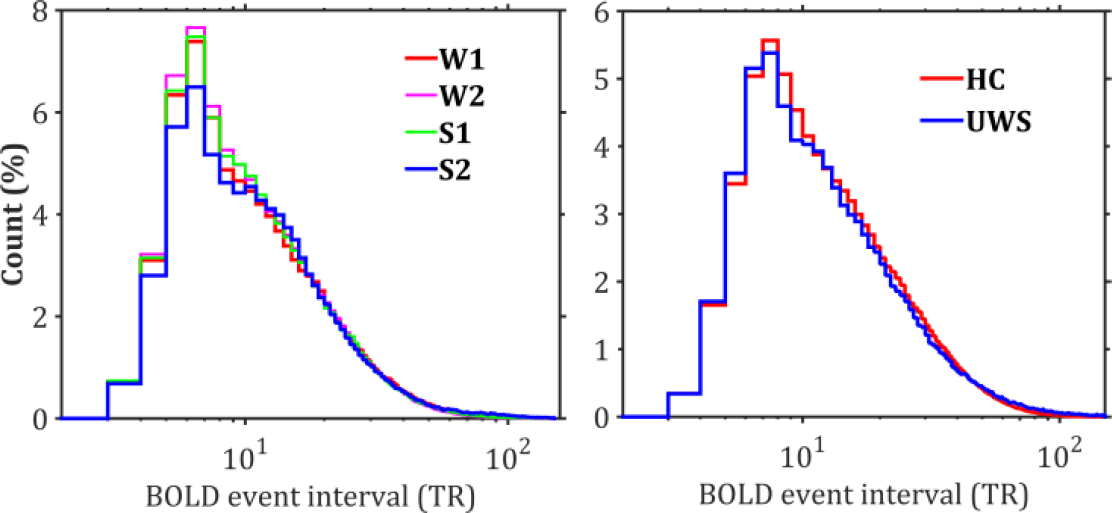
Group level frequency distribution: of temporal intervals between adjacent point process events. Left: Propofol dataset; Riglit: UWS dataset.

**Figure 3.**
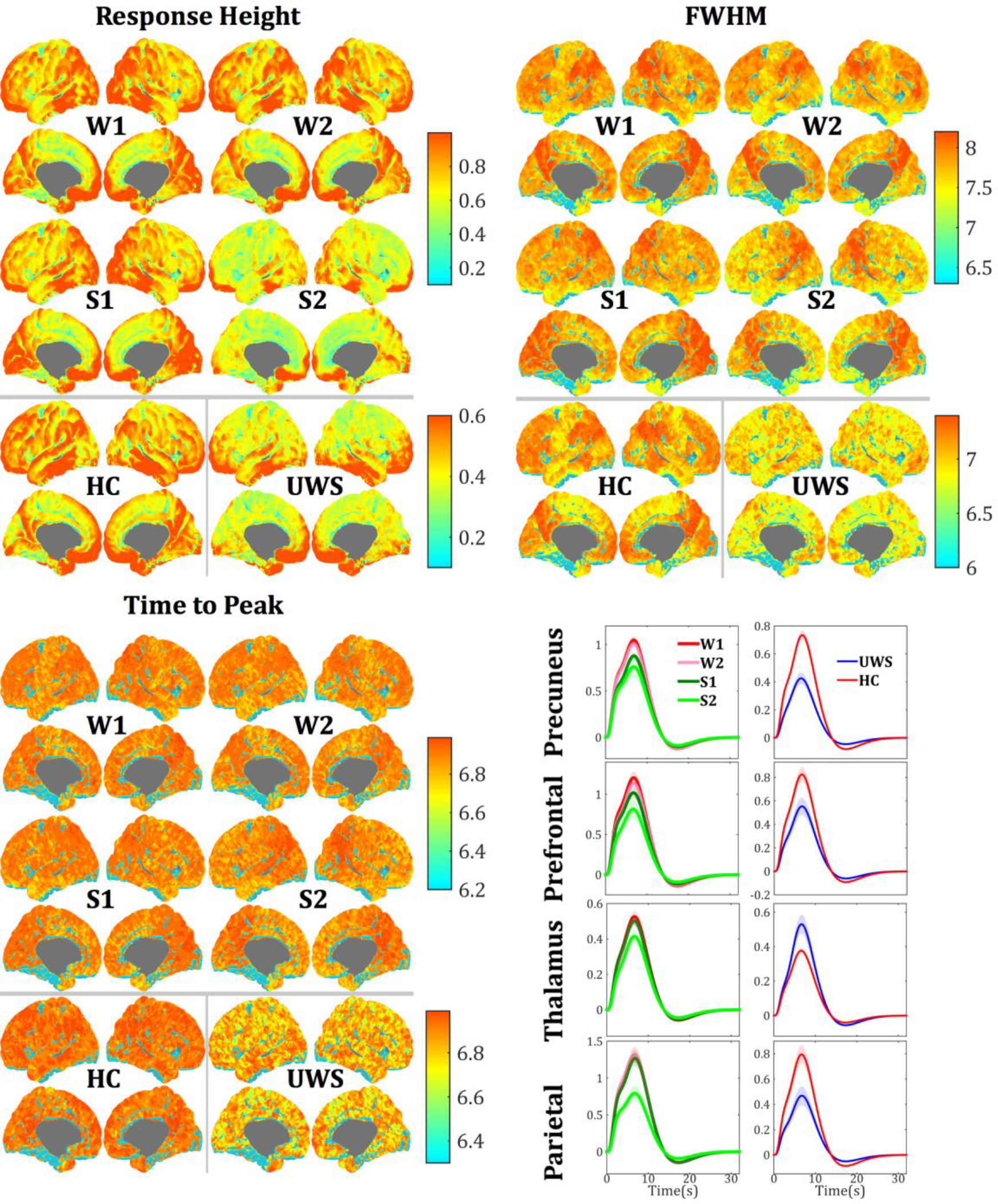
Median maps of HRF parameters (response height, FWHM, time to peak), estimated by canon2dd model. Different scales are used for each dataset and HRF parameter, while the same scale is used for the same HRF parameter and dataset. For better comparison, the maximum value in colormap is not the maximum value of HRF parameters. The lower right panel reports the group averaged HRF as percent signal change with its standard error in four representative locations (precuneus, MNI coordinates [−3, −60, 27]; thalamus, MNI coordinates [−9, −9, 3]: prefrontal, MNI coordinates [51,39,13]; parietal, MNI coordinates [−45, −57, 51]). The left column depicts results from the Propofol dataset, the right one from the UWS dataset.

### Group HRF differences and Conjunction

Statistical maps of HRF parameters reveal that HRF shape parameters are modulated by Propofol anesthesia (Figure 4). The linear covariation between HRF and the level of consciousness is only evident in specific brain area. Such phenomenon mostly occurs in the response height: frontal lobe (middle/medial/inferior/superior frontal gyrus), anterior cingulate, inferior/superior parietal lobule, precuneus, posterior cingulate, supramarginal gyrus, angular gyrus, precentral/postcentral gyrus (primary somatosensory/motor cortex), supplementary motor area, superior/middle temporal gyrus, insula, parahippocampus, hippocampus.

**Figure 4.**
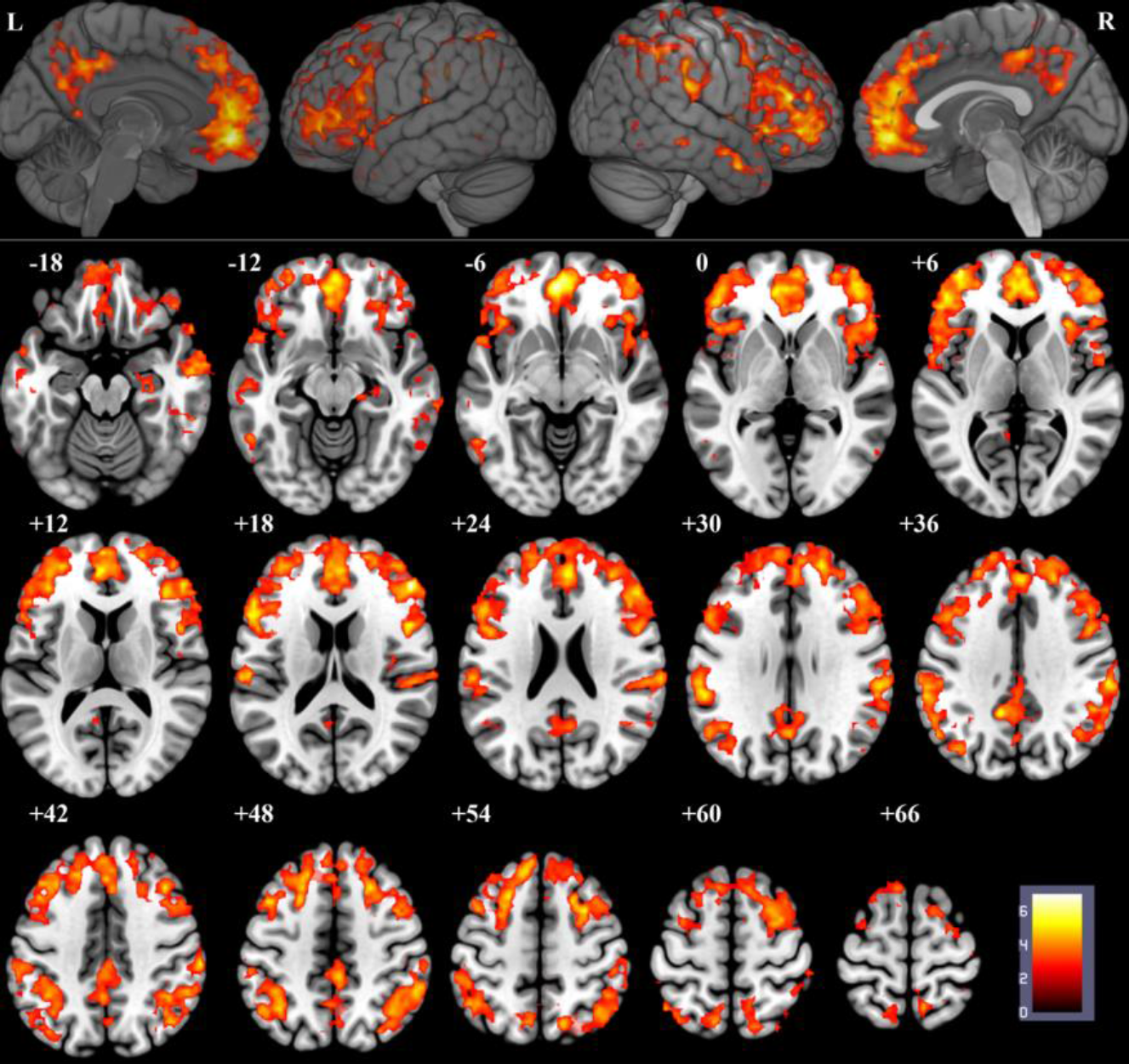
Linear correlation between response lieight and four levels of consciousness (Wl, W2, Sf, S2), p<0.05, topo FDR Correction.

Group pairwise differences between the two most extreme cases (W1 and S2) show a similar spatial distribution (Figure 5).

**Figure 5.**
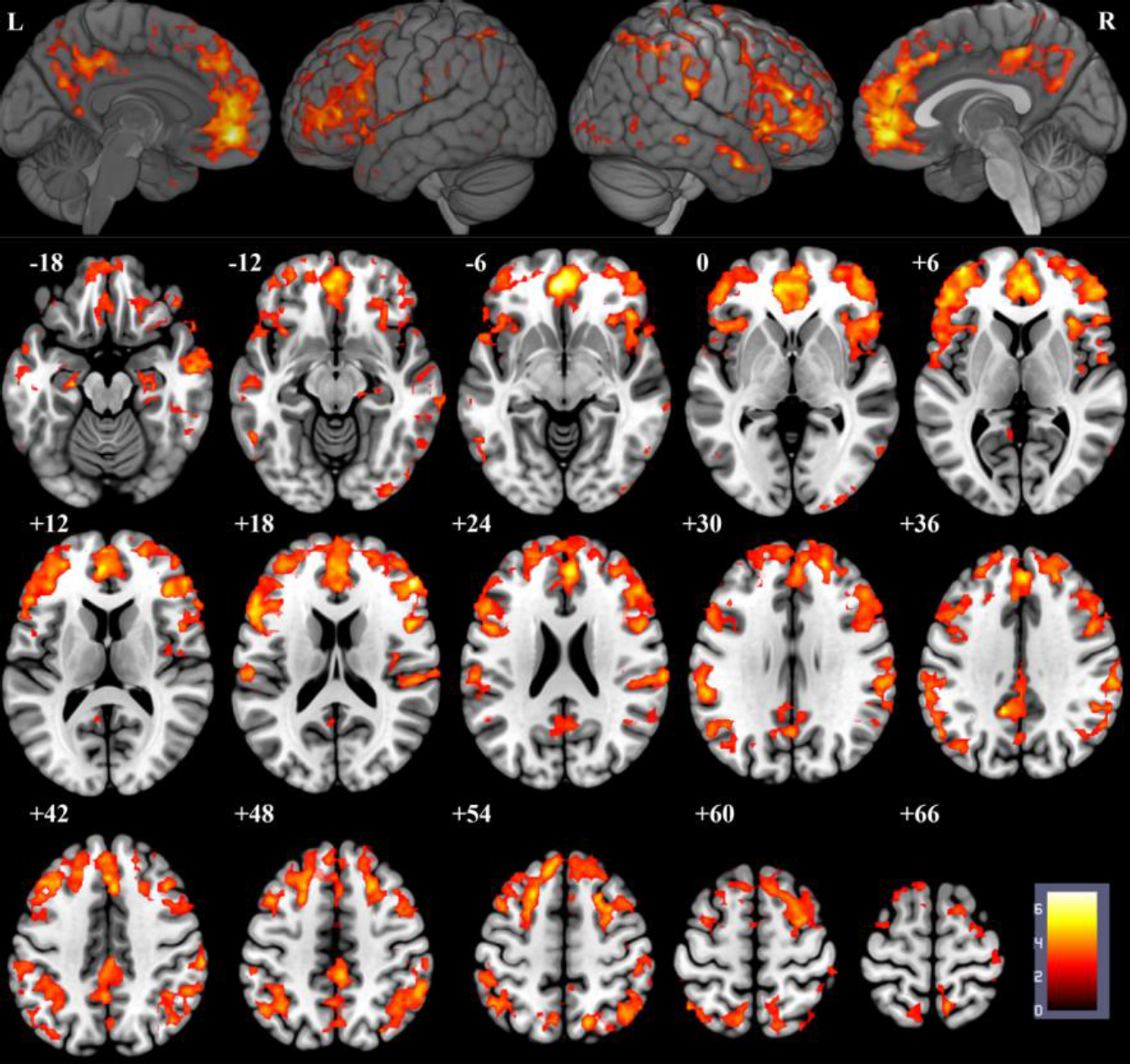
Response height differences between W1 and S2, p<0.05, topo FDR correction.

A linear relationship can also be found in FWHM in the frontal gyrus (medial/middle/inferior/superior) and anterior cingulate (Figure 6.A). No linear relationship was found in temporal latency of hemodynamic response.

**Figure 6.**
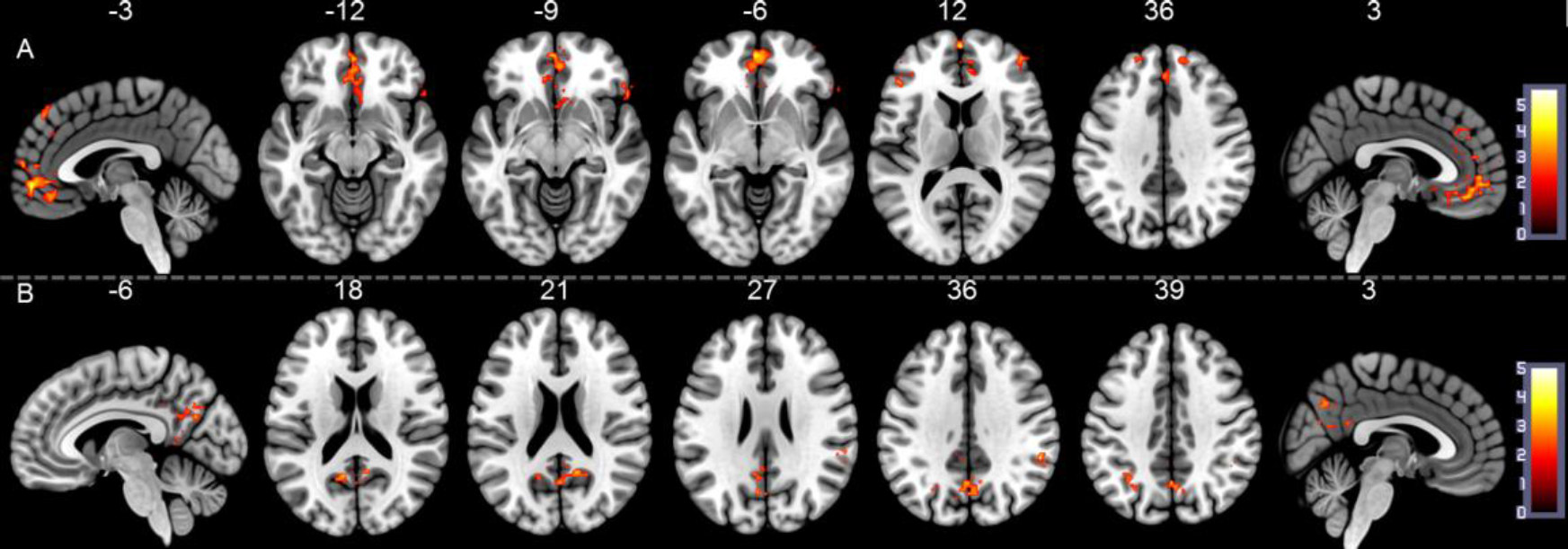
(A) linear correlation between FWHM and four levels of consciousness (wl, w2, s1, s2), p<0.05, topo FDR correction. (B) Group differences of FWHM between healthy controls and UWS patients, p<0.05, topo FDR correction.

The UWS subjects show lower spontaneous response height than control subjects in precuneus, posterior cingulate, cuneus, inferior parietal lobule, supramarginal gyrus, and angular gyrus, but higher response height in put amen, thalamus, parahippocampus gyrus, fusiform, and anterior cingulate, While the sub-regions of frontal gyrus exhibit contrary different patterns in spontaneous hemodynamic response height between UWS and control group (Figure 7). In contrast to the UWSgroup, the control subjects show broader response width (FWHM) in precuneus, posterior cingulate, inferior parietal lobule, supramarginal gyrus (Figure 6.B). No group difference is found in the time to peak of hemodynamic response.

**Figure 7.**
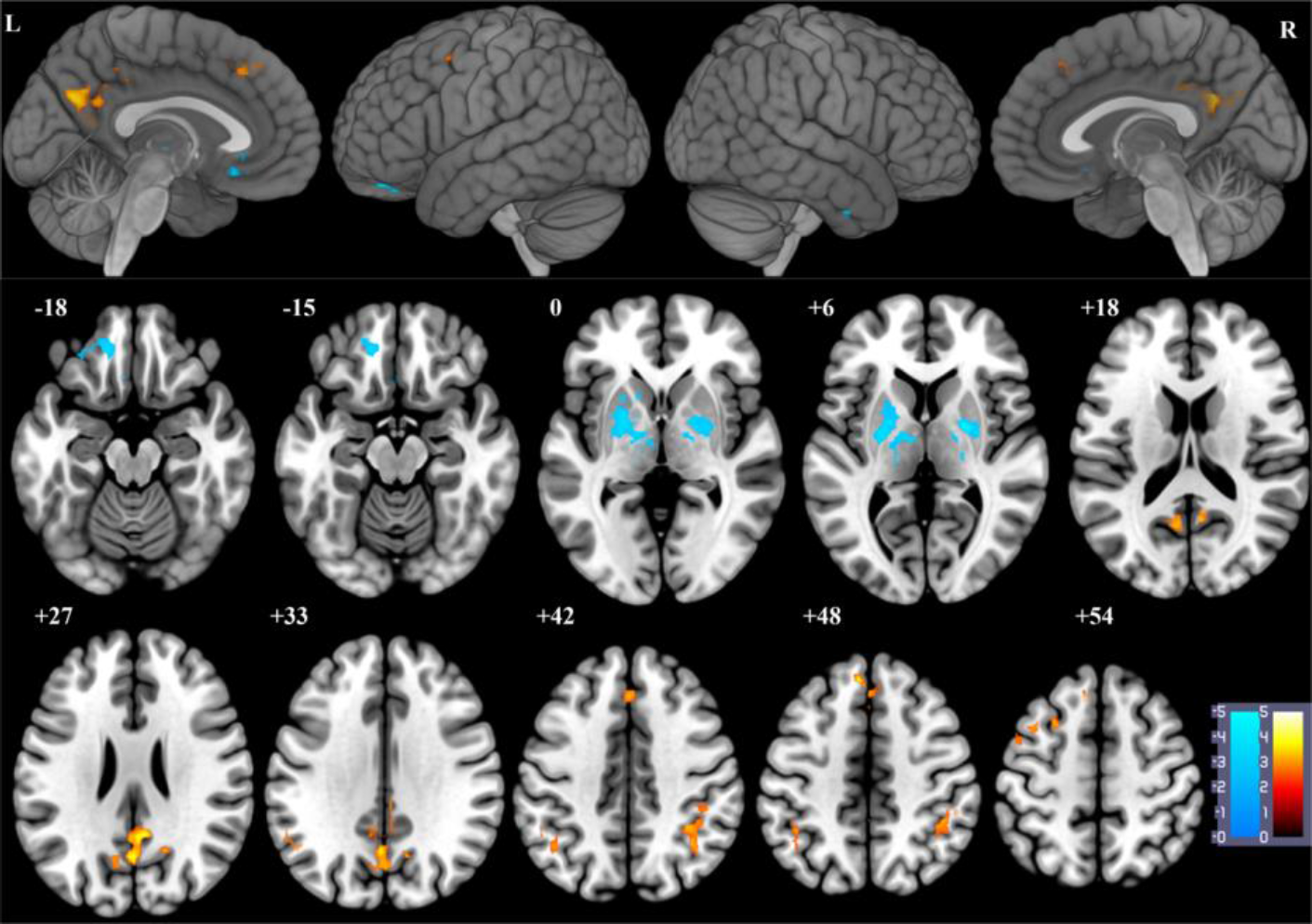
Response height differences between healthy controls and UWS patients, p<0.05, topo FDR correction.

A conjunction analysis for W1 minus S2 and Control minus UWS contrasts yields a significant cluster of higher spontaneous HRF response height in precuneus and posterior cingulate (Figure 8).

**Figure 8.**
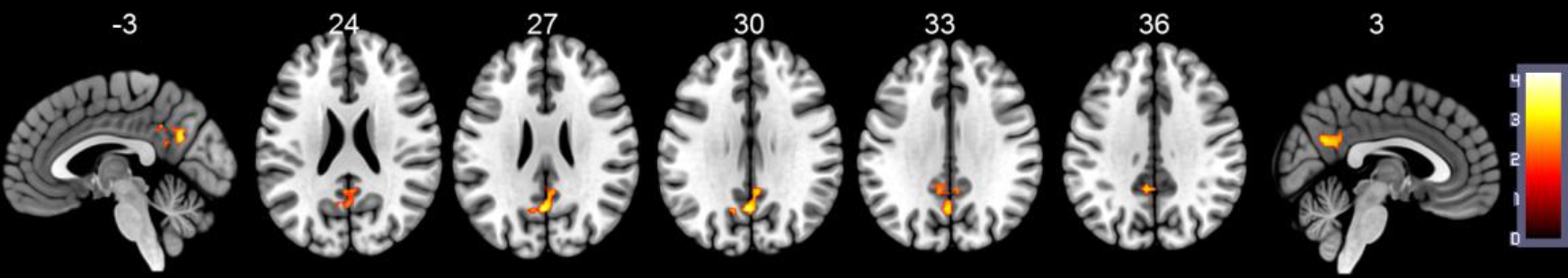
Conjunction map of W1 minus S2 and healthy control minus UWS patients in response height differences, p<0.05, topo FDR correction.

## Discussion

We investigated the modulations induced by changing levels of consciousness in resting state hemodynamic response based on BOLD fMRI signals in healthy control with Propofol anesthesia and UWS patients. This allowed us to explore the dissociation between wakefulness and awareness, present in UWS but not in anesthesia, from the point of view of a measure that is relevant both to the dynamics (local and integrated), and to the local metabolism.

The hemodynamic response showed both similarities and differences in pharmacological and pathologically induced loss of consciousness. We found that a modulation of hemodynamic response in precuneus and posterior cingulate is a common principle underlying loss of consciousness induced by Propofol anesthesia and UWS, which is consistent with previous PET and functional connectivity MRI studies (Fiset et al., 2005; Laureys, 2005). The hemodynamic response in frontoparietal networks is clearly altered with Propofol anesthesia, but changes in subcortical regions like the thalamus are not as evident. However, the spontaneous thalamic hemodynamic response exhibits distinctly different characteristic between healthy controls and UWS patients.

It is well established that most anesthetic agents decrease glucose metabolism in a dose-dependent manner with variable effect on glucose metabolism rate (GMR) and cerebral blood flow (CBF) (Alkire and Miller, 2005). While the BOLD-fMRI signal reflect the complex interactions between cerebral metabolic rate of oxygen, cerebral blood flow and volume (Ogawa et al., 1990). Therefore, the changes that occur in the brain metabolism and CBF caused by neural activity could be accompanied by concurrent changes in BOLD effect in a predictable way. Most fMRI studies employ active or passive paradigm to detect activation in sensory or cognitive system (Laureys and Schiff, 2012). However the quantifiable nature of relation between cerebral metabolism and resting state BOLD HRF has been left largely unexplored. Many studies focused on regional GMR and CBF with PET to measure anesthesia or pathology induced unconsciousness (Alkire and Miller, 2005; Laureys, 2005). In the unconscious state, the consistent regions that show relative decreases of metabolism and blood flow are in the frontoparietal networks, DMN, as well as the thalamus (Alkire and Miller, 2005; Laureys et al.. 2004; Nakayama et al., 2006). These regions are involved in conscious processing. The loss of consciousness suggests a disturbance in the optimal balance between segregation and integration among these regions and the connected regions. Such as DMN is considered to be involved in stimulus-independent thought, mind-wandering and self-consciousness (Raichle, 2015). The posterior cingulate cortex (PCC) and the medial precuneus are prominent features of the DMN, they act as a core hub role in integrating information across functionally segregated brain regions, display high resting metabolically active. PCC has dense structural connectivity to widespread brain regions, is involved in internally directed thought (Leech et al., 2012), and engaged in continuous information gathering and representation of the self and external world with interconnected precuneus and medial prefrontal cortices (Gusnard et al., 2001). The precuneus is one of the first regions of the brain to resume activity when regaining consciousness from a UWS, and together with the adjacent PCC are the regions that differentiate patients in minimally conscious states from those in UWS (Laureys et al., 2004). Therefore, the consistent evidence of decreased hemodynamic response in precuneus and posterior cingulate speaks to deficits in cognitive functioning and information integration possibly leading to a breakdown of consciousness.

A BOLD-fMRI study on brain activation by auditory word stimulus during Propofol sedation suggested that the superior and middle temporal gyrus and the inferior parietal lobule are. associated with the formation of both implicit and explicit memories (Quan et al., 2013). The present results exhibit that Propofol effect on explicit and implicit memory can be discovered from baseline hemodynamic response. The thalamus plays an important role in normal arousal regulation (Schiff, 2008), thus is always the common locus of action of brain injury in UWS and of general anesthetics. Its anatomy and physiology imply a central role in consciousness (Ward, 2011). Certainly the thalamic activity is identified as a key target of anesthetic effects on consciousness. However, thalamic depression may depend on the agent studied and the degree of sedation. It has been suggested that cortical cells are more sensitive to the effect of Propofol than sub-cortical elements (Alkire and Miller, 2005; Sun et al., 2008). Our finding confirms such phenomenon, that is spontaneous hemodynamic response in frontal lobe is significant correlated with Propofol anesthesia, but such effect cannot be identified in thalamus due to the cluster size cannot achieved significant threshold. Nonetheless, significant positive linear correlation with hemodynamic response height in thalamus could be found with small volume correction, and some negative linear correlation in the part of thalamus (p<0.05, uncorrected, Figure 9), while only higher hemodynamic response exhibited in UWS compared with healthy control. However, it does not imply the thalamic activity failed to be suppressed by Propofol. The Propofol effect may mediated the thalamocortical interaction (Alkire et al., 2008). Indeed, the thalamocortical connectivity in default network and bilateral executive-control networks were found to be correlated with Propofol-induced decrease of consciousness in previous study on this dataset (Boveroux et al., 2010). Meanwhile, an increase in FC between the thalamus and the auditory cortex, insular cortex, primary somatosensory cortex, primary motor cortex, and supplementary motor area, was observed with mild sedation (Guldenmund et al., 2013). In addition, the hyperactivity in thalamus is consistent with limbic hyperconnectivity in UWS shown in a recent study (Di Perri et al., 2013). This may reflect the correlation between alpha activity increases and spontaneous hemodynamic response (Brown et al.. 2010; Wu and Marinazzo, 2015). The evident decrease of spontaneous hemodynamic responses in decreased states of consciousness offers a strong empirical support that thalamus is at the heart of the neurobiology of consciousness.

**Figure 9.**
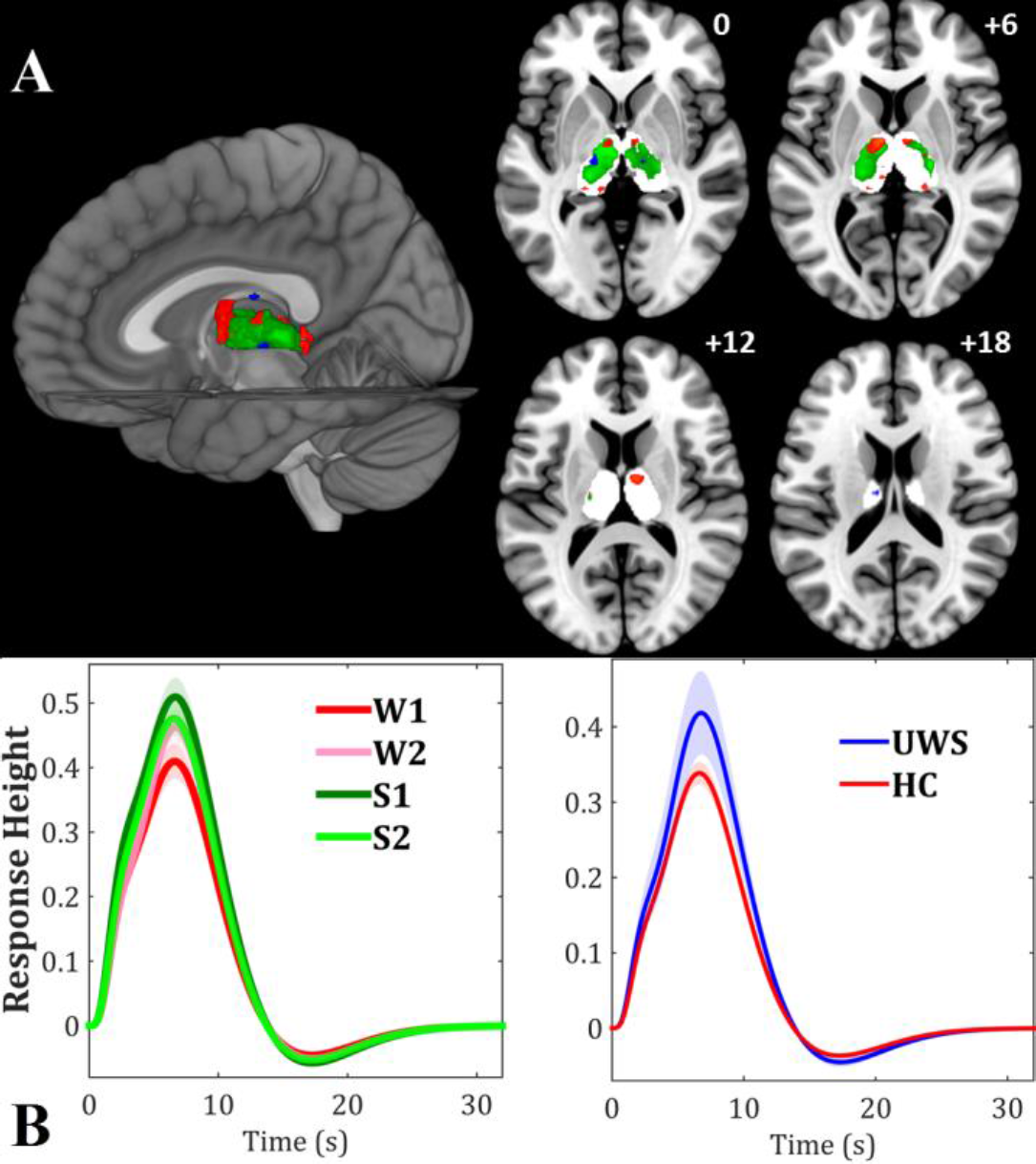
(A): W1 minus S2 (positive: red, negative: blue) and healthy control minus UWS patients (negative: green) in response height differences, p<0.05, uncorrected. (B): Group averaged HRF with its standard error in a thalamus (white shadow mask) voxel (MNI coordinates: [−15, −18, 0])

It’s worth to mention that the possible relationship between resting state FC and brain activity reflect by hemodynamic response in the pathological or pharmacological coma remains unclear. Hyperconnectivity or hypoconnectivity only indicate changes of synchronous cortical activity among brain areas, which may be induced by only one region or both of them, further analysis on hemodynamic response in each area could clearly explain the link between brain connectivity and activity (Tomasi and Volkow, 2018). The effects of hemodynamic variability on resting state FC have been investigated (Rangaprakash et al., 2018), as well as their implications for posttraumatic stress disorder (PTSD) (Rangaprakash et al., 2017) and autism (Yan et al., 2018).

### Methodological considerations

Stimulus-evoked BOLD fMRI is one of the most conventional paradigms to investigate anesthetic influence on neuronal activation. These studies rely on the response of specific brain regions to a given task. On the other hand unexpected or unspecific widespread BOLD signal changes across the brain are frequently observed. Finding a balance between complexity and applicability is always a challenge for task stimuli. Conversely, resting state fMRI is a paradigm-free fMRI approach, and allows investigating the baseline activity of the whole brain. Resting state functional connectivity analysis is the most widely used technique in anesthesia and pathology resting state fMRI studies. Due to relatively poor temporal resolution, few studies focus on exploring the spontaneous event dynamics of anesthesia. As shown in previous works, an efficient way to reveal this spontaneous activity at cortical level from resting state BOLD signal could be through point process analysis, under the hypothesis that important features of brain dynamics at rest can be captured from relatively large fluctuations in BOLD amplitude (Tagliazucchi et al., 2012; Wu et al., 2013). A recent study on PCC related co-activations patterns by point process analysis have shown differentiate region-specific Propofol-induced reductions on brain functional network organization (Amico et al., 2014).

As discussed in (Di et al., 2008), due. to uncontrolled head movements during scanning, the patients with UWS may show more motion artifacts than in collaborative healthy subjects. To avoid motion-related artifacts contribution to point process, 3Ddespike and aCompCor were combined to attenuate motion artifacts, to minimize the framewise relationship between head motion and signal change (Muschelli et al., 2014). To further minimize the motion artifact influence on HRF shape retrieved from point process, data scrubbing is performed (Power et al., 2012), and mean FD power of each subject is included as a covariate for further statistical analysis (Van Dijk et al., 2012). This procedure indicates that our finding is unlikely to be due to motion artifact.

Physiological (cardiac, pulmonary) contributors of BOLD signal, and sources of its variability, should be taken into account, especially in resting state fMRI studies. These also influence the detection of spontaneous events and the retrieved HRF (Wu and Marinazzo, 2016). The datasets used here did not contain explicit physiological measurements, nonetheless the use of aCompCor represent the next best solution. The GLM based HRF retrieving will benefit from heterogeneous distribution of temporal interval between these point process events, i.e. temporal ‘jitter’ variation in event onset times (Buckner, 1998), while its power is relative to violation the assumption of linear additively (Boynton et al., 1996). As shown in a simulation study, there is more bias in the estimates of FWHM and time to peak than response height (Lindquist et al., 2009). These evidences could partially explain less group level finding discovered in FWHM and time to peak. Meanwhile;, our findings also suggest that hemodynamic response height shows the most discriminative power than other HRF parameters. Additionally, the coupling between neural activity and the vascular response is significant in determining the amplitude and spatial resolution of the BOLD signal (Logothetis and Wandell, 2004). Regions of sparse vascularization are likely to have low efficacy of this coupling and weak or absent BOLD response. This may be related to higher hemodynamic response height found in the cortical surface.

To conclude, our results show that neurovascular coupling in different states of consciousness could be tracked by modulated spontaneous hemodynamic response derived from resting state fMRI. These results demonstrate the feasibility of resting state HRF for the study of the brain at rest, revealing comprehensive and complementary information to further decode anesthetic and pathological brain function, especially for clinical populations and conditions not suitable for PET imaging.

## Acknowledgements

G-R.W was supported by the National Natural Science Foundation of China (Grant No. 61876156)

